# Conserved epigenetic programming and enhanced heme metabolism drive memory B cell reactivation

**DOI:** 10.1101/2021.01.20.427446

**Authors:** Madeline J. Price, Christopher D. Scharer, Anna K. Kania, Troy D. Randall, Jeremy M. Boss

## Abstract

Memory B cells (MBCs) have enhanced capabilities to differentiate to plasma cells and generate a rapid burst of antibodies upon secondary stimulation. To determine if MBCs harbor an epigenetic landscape that contributes to increased differentiation potential, we derived the chromatin accessibility and transcriptomes of influenza-specific IgM and IgG MBCs compared to naïve cells. MBCs possessed an accessible chromatin architecture surrounding plasma cell specific genes, as well as altered expression of transcription factors and genes encoding cell cycle, chemotaxis, and signal transduction processes. Intriguingly, this MBC signature was conserved between humans and mice. MBCs of both species possessed a heightened heme signature compared to naïve cells. Differentiation in the presence of hemin enhanced oxidative phosphorylation metabolism and MBC differentiation into antibody secreting plasma cells. Thus, these data define conserved MBC transcriptional and epigenetic signatures that include a central role for heme and multiple other pathways in augmenting MBC reactivation potential.

**Key Points:**

- Influenza-specific memory B cells have accessible chromatin structure.
- Human and mouse memory B cells upregulate heme metabolic pathways.
- Heme enhances PC differentiation and augments mitochondrial metabolism in ex vivo.

## INTRODUCTION

Long-lived humoral immunity is realized through the joint functions of memory B cells (MBCs) and plasma cells (PCs). Naïve B cells (nBs) and MBCs have similar fates upon T-dependent stimulation although MBCs respond more quickly and with greater magnitude to antigenic challenge (1, 2). For the most part, MBCs tend to be affinity matured with class-switched BCRs (3–5) and gain unique surface markers, such as CD73, PD-L2, and CD80 (6, 7). It is now also appreciated that IgM^+^ MBCs are also key players in memory responses (8, 9). For class-switched MBCs, enhanced immune responses have been attributed in part to the signaling capacity of the IgG versus IgM BCR (4). However, naïve cells genetically engineered to express the IgG BCR respond similarly to IgM^+^ nBs (10). Together, these data suggest that alternative mechanisms function to promote heightened MBC functional responses.

One possibility is that MBCs have acquired epigenetic programming that distinguishes them from their naïve precursors. Evidence supporting this hypothesis includes the role of the monocytic leukemia zinc finger protein (MOZ), a histone acetyltransferase that targets H3K9 and is required for high affinity class-switched MBCs (11). Human MBCs harbor an unique DNA methylation profile compared to naïve, germinal center (GC) B cells, and PCs (12, 13). In a similar manner, memory CD8 T cells maintain a primed chromatin accessibility architecture at key effector genes (14). Changes in the levels of transcription factors also correlate with MBC properties. For example, Krüppel-like factor 4 (KLF4) restricts the proliferative response and is decreased in expression in MBCs compared to naïve. Similarly, down regulation of BACH2, a repressor of BLIMP-1 and the PC fate, has been proposed as one mechanism for enhanced MBC differentiation potential (10). In contrast, upregulation of ZBTB32 restrains MBC secondary responses (15). Despite this understanding, the molecular programming of MBC subsets remains incomplete.

Here, we define the transcriptional and chromatin accessibility profiles of IgM and IgG MBC subsets compared to nBs. We found that MBCs harbor distinct epigenetic programming from nBs that is conserved between the MBCs of mice and humans. This programming includes the priming of accessible chromatin surrounding PC specific genes, some of which are BLIMP-1 targets. Moreover, we found that heme biosynthesis was a conserved key feature of the MBC state, in which augmented differentiation to PCs was observed in the presence of the iron-containing complex hemin. In summary, our findings describe unique epigenetic parameters and features of MBC programming that allow these cells to respond more quickly and vigorously to secondary immune challenges.

## MATERIALS AND METHODS

### Data availability

Sequencing data generated in this study have been deposited in the NCBI Gene Expression Omnibus (https://www.ncbi.nlm.nih.gov/geo/) and are available under accession GSE164173. Code used to analyze the data and generate the figures is available from http://github.com/cdschar and upon request.

### Mouse influenza infection

C57BL/6J wild-type mice were obtained from The Jackson Laboratory (000664). All experiments were performed with a mix of male and female mice that were between 8-10 weeks of age. All animals were housed in specific pathogen-free cages by the Emory Division of Animal Resources and all protocols were approved by the Emory Institutional Animal Care and Use Committee (IACUC). Both male and female mice that were between 8-10 weeks of age were infected with influenza A/PR8/34 (PR8) at 15,000 virus forming units (vfu) per dose. Greater than 35 days post infection, mice were sacrificed or challenged with a 0.1 median lethal dose (LD_50_) of influenza A/HK-X31 (X31). For all infections, mice were anesthetized with vaporized isofluorane and 30 μl of PR8 or 100 μl of X31 were administered intranasally.

### Mouse B cell isolation

Splenic cell suspensions were made by mechanically forcing spleens through a 40 μm filter and lysing red blood cells with ACK lysis buffer (0.15 M NH_4_Cl, 10 mM KHCO_3_, 0.1 mM EDTA) for 3 mins before quenching the reaction with four volumes of complete RPMI media: RPMI 1640 (Corning Cellgro, 50-020-PC), 10% heat-inactivated FBS (Sigma-Aldrich), 1% MEM non-essential amino acids (Sigma, ENBF3930-01), 100 μM sodium pyruvate (Sigma, RNBF6686), 10 mM HEPES (HyClone, SH30237), 1x Penicillin-Streptomycin-Glutamine (Gibco, 10378016), 0.0035% β-mercaptoethanol (Sigma-Aldrich). Naive B cells were isolated using negative selection for CD43 (Miltenyi, 130-090-862) following the manufacturer’s protocol. Purity was confirmed by flow cytometry. B cell antigen tetramers were described previously (16). Influenza nucleoprotein (NP)-specific memory B cells were enriched using B cell tetramer pull-downs. Cells were stained with 1:100 NP-phycoerythrin (PE) for 30 mins, washed, and incubated with anti-PE microbeads (Miltenyi, 130-105-639) before positive selection with magnetic columns. Memory B cells for ex vivo cultures were enriched using the Memory B cell Isolation Kit, Mouse (Miltenyi, 130-095-838) following manufacturer’s protocol.

### Mouse ex vivo T-dependent stimulation

Naïve B cells and total enriched memory B cells were isolated as described above, resuspended at 0.5 x 10^6^ cells per ml, and stimulated with 500 ng/ml soluble CD40L (R&D Systems, 8230-CL), 10ng/ml IL-4 (R&D Systems, 404-ML), and 10 ng/ml IL-5 in complete RPMI. Cultures were fed on each subsequent day with 10 ng/ml IL-4 and 10 ng/ml IL-5 until harvest at the indicated time points. For hemin experiments, 60 μM hemin (Sigma) or vehicle (NH_4_OH) was added to cultures on day 1.

### Human B cell isolation

Peripheral blood mononuclear cells (PBMCs) were obtained from deidentified healthy donors according to approved protocols from the Emory Institutional Review Board (IRB). For each isolation, 40 ml of blood 1:2 with PBS, followed by layering over a 50% volume of Ficoll (17-1440-02), and centrifuged at 1850 rpm for 30 mins with minimal acceleration and deceleration. The lymphocyte layer was removed, and human memory B cells enriched using the human memory B cell kit (Miltenyi, 130-093-546) following the manufacturer’s protocol.

### Human MBC ex vivo stimulation

Enriched human memory B cells were cultured at 0.2 x 10^6^ cells per ml in complete RPMI supplemented with 1 mg/ml R848 (Invivogen), 1 mg/ml soluble CD40L (Fitzgerald), 10 ng/ml IL-21, 50 U/ml IL-2, 10 ng/ml BAFF (Peprotech), and 0.8 μl/ml human insulin for 6 days. On day 2 of the culture, vehicle or 60 μM hemin was added as indicated.

### Flow cytometry

Staining panels for mouse included anti-Fc (anti-CD16/CD32) (Tonbo Biosciences, 2.4G2), anti-CD11b (Tonbo Biosciences, M1/70), anti-CD11c (Tonbo Biosciences, N418), anti-F4/80 (Biolegend, BM8), and Thy1/2 (Biolegend, 30-H12) conjugated to APC-Cy7 each at a concentration of 0.25 μg per 1 x 10^6^ cells to remove dendritic cells, macrophages, and T cells. The following stains and antibody-fluorophore combinations were used to assess cellular phenotype: anti-B220-PE-Cy7 or -AlexaFluor700 (Tonbo Biosciences, RA3-6B2) at 0.5 μl per 1 x 10^6^ cells; anti-CD138-APC, or -BV711 (BD, 281-2) at 0.125 μl per 1 x 10^6^ cells; anti-GL7-PerCp-Cy5.5, -PE, or -e450 (Biolegend, GL7) at 0.25 μl per 1 x 10^6^ cells; anti-CD38-Pacific Blue (Biolegend, 90); anti-IgD-BV711 (BD, 11-26c.2a); anti-IgM-FITC (Invitrogen, II/41); anti-IgG-PerCp-Cy5.5 (Biolegend, Poly4053); Zombie Yellow (BV570) (Biolegend, 77168), and CTV (Life Technologies, C34557). CTV was used at 10 μM per 1 x 10^7^ cells/ml. Human staining panels included anti-CD3-Alexa 700 (Invitrogen, UCHT1), anti-CD19-APC (Biolegend, SJC25C1), anti-CD27-BV786 (BD L128), anti-IgD-APC-Cy7 (Biolegend, IA6-2), anti-CD38-PE-Cy7 (Biolegend, HIT2), and anti-CD138-PerCp-Cy5.5 (BD, MI15) or anti-CD138-Alexa700 (Biolegend, MI15), all used at 5 μl per 1 x 10^6^ cells. Intracellular HO-1 staining for human and mouse was performed using the Invitrogen FIX & PERM Kit (GAS003) using HO-1 PE at 1:100 dilution (Abcam, ab83214) and PE IgG2b isotype (Abcam, ab91532) for gating. Staining panels included fluorescence minus one (FMO) controls to ensure that correct compensation and gating were applied. BD LSR II or BD LSR Fortessa X-20 was used for analysis and a FACSAria II was used for sorting (BD Biosciences). All flow cytometry data were analyzed with FlowJo v10.

### RNA-seq

Naïve B cells were isolated from the draining lymph node of age-matched naïve mice and sorted as follows: Live CD11b^-^F4/80^-^Thy1.2^-^B220^+^CD38^+^GL7^-^IgM^+^. Memory B cells were isolated from 4 pooled memory mice and sorted as follows: Live CD11b^-^F4/80^-^Thy1.2^-^B220^+^CD38^+^GL7^-^NP-PE^+^NP-APC^+^IgM^+^ or IgG^+^. 500 cells were isolated via FACS into 300 μl RLT buffer (Qiagen) containing 1% pME. At the time of RNA isolation, 3 μl of a 1:400,000 dilution of ERCC synthetic mRNAs (ThermoFisher) were added and total RNA was isolated using a Quick-RNA MicroPrep Kit (Zymo Research, 11-328M) and all subsequent RNA used for cDNA synthesis using the SMART-seq v4 kit (Takara) and 10 cycles of PCR. To generate final libraries 200 μg of cDNA was used as input for the NexteraXT DNA Library kit (Illumina) using 10 additional cycles of PCR. An Agilent Bioanalyzer was used to quality control check each library. Libraries were pooled at an equimolar ratio and sequenced on a NextSeq 500 (Illumina) using 75 base pair paired-end chemistry.

### RNA-seq analysis

Raw sequencing reads were mapped to the mm10 genome using STAR (17). Gene counts were determined with the Bioconductor package GenomicAlignments (18) using the mm10 transcriptome database. The Bioconductor package edgeR (19) was used to determine differentially expressed genes (false discovery rate (FDR) ≤ 0.05, absolute log_2_FC ≥ 1). All differentially expressed genes are listed in **Supplemental Table 1**. Hierarchical clustering boostrapping analysis was performed using the pvclust v2.2-0 R package with 1000 replications. GO analysis was performed with DAVID (20). Preranked Gene Set Enrichment Analysis (21) was performed on all detected transcripts ranked by multiplying the sign of fold change by the -log_10_ of the edgeR *P* value. Normalized mRNA content was calculated as previously described (22). To compare gene expression between species the ENTREZ ID for both human and mouse was used as input for the NCBI HomoloGene database (23). Only unique matches were used for downstream analysis resulting in 8,639 unique genes that were detected in both species.

### ATAC-seq

Naïve and memory B cells were sorted as defined above. At least 1,000 cells for each cell type were isolated via FACS for ATAC-seq. Libraries were generated as described in detail previously (24). Briefly, cells were centrifuged at 500 x g for 10 min at 4°C. The supernatant was removed with a pipette and cells resuspended in 25 μl Tn5 Reaction Buffer (2.5 μl Tn5, 12.5 μl 2x TD buffer, 5 μl molecular grade H_2_O, 2.5 μl 1 % Tween-20, and 2.5 μl 0.2% Digitonin) and incubated at 37°C for 1 hr. Cells were lysed by adding 2 μl 10 mg/ml Proteinase-K and 23 μl Lysis Buffer (326 mM NaCl, 109 mM EDTA, 0.63% SDS) and incubated for 30 min at 40°C. Large molecular weight DNA was removed by addition of 0.7x volumes of AMPureXP SPRI-beads (Invitrogen Agencourt, A63880) and low molecular weight DNA isolated for downstream analysis using a 1.2x volumes AMPureXP SPRI-beads and eluted in 15 μl EB buffer (Qiagen). ATAC-seq libraries were amplified using 5 μl each of the i5 and i7 Nextera Indexing primers and 25 μl of 2x HiFi HotStart ReadyMix (Roche Diagnostics, KK2601). Final libraries were purified using 1x volumes of AMPureXP SPRI-beads and sequence using 75bp paired-end reads on an Illumina NextSeq500.

### ATAC-seq analysis

Raw sequencing reads were mapped to the mm10 genome using Bowtie (25). Peaks were called using MACS2 (26) and annotated to the nearest transcription start site using HOMER (27). Data were normalized to reads per peak per million (rppm) as previously described (28) and all samples demonstrated a fraction of reads in peaks (FRiP) score of 21% or greater. The coverage for all peaks was annotated using the GenomicRanges (18) package and edgeR (19) waw used to determine differential accessibility between samples (FDR ≤ 0.05, absolute log_2_FC ≥ 1). All DAR are listed in **Supplemental Table 2**. Motif analysis was performed with the HOMER program findMotifsGenome.pl using randomly generated genomic sequences as background. PageRank analysis (29) was performed using the default parameters and transcription factors with a PageRank score >0.001 in at least one sample group of the indicated comparison were included for all downstream analysis.

### Quantitative RT-PCR

Cells were lysed in 300 μl RLT buffer (Qiagen) containing 1% pME and total RNA was isolated using the Quick-RNA MicroPrep Kit (Zymo Research, 11-328M). All subsequent RNA was used for reverse transcription using random hexamer and oligo dT primers with SuperScriptII (ThermoFisher, 18064022). cDNA was diluted 1:5 and gene expression quantitated by real-time PCR using SYBR Green incorporation on a Bio-Rad iCycler. Gene expression was quantitated as a percentage of 18S rRNA. All primer sets are listed in **Supplemental Table 3**.

### ELISPOTS

For ELISPOT assays, multiscreen HA plates (Millipore, MAHAS4510) were coated with purified NP at 1 μg/ml in PBS overnight at 4°C. Plates were washed four times with PBS and blocked with complete RPMI. Single-cell suspensions were washed, diluted in complete RPMI, and cultured on coated plates overnight in a 37°C incubator with 5% CO_2_. Cells were aspirated, and plates were washed with 0.2% Tween-20 in PBS. Bound IgG was detected using alkaline phosphatase–conjugated goat anti–mouse kappa (Southern Biotech, diluted 1:1000) in 0.5% BSA, 0.05% Tween-20 in PBS for 1 hr at 37°C. Plates were washed with 0.2% Tween-20 in PBS and developed with BCIP/NBT (Moss Substrates) substrate for 1 hr. ImmunoSpot S6 ULTIMATE Analyzer (Cellular Technology Limited; CTL) and quantified using ImmunoSpot software v5.0.9.21 using the Basic Count feature with the following parameters: Sensitivity: 165; Background balance: 162; Diffuse Processing: Largest; Spot size range: 0.0201 mm^2^ – 9.626 mm^2^; Spot Separation: 1.

### Metabolism assays

A FluxPack cartridge was hydrated for 12 hours prior to each assay with 200 μl dH_2_O at 37°C in a non-CO_2_ incubator. Approximately 1 hr prior to each assay, dH_2_O was replaced with pre-warmed Seahorse calibrant solution (Agilent, 103059-000). Cell cultures were washed in Seahorse XF Assay media supplemented with 1 mM sodium pyruvate, 2 mM L-glutamine, and 5.5 mM glucose. Cells concentrations were determined by flow cytometry using AccuCheck counting beads (Invitrogen, PCB100). CellTak (Corning, 354420) was diluted in sterile 1xPBS to a final concentration of 22.4 μg/ml and 25 μl added to each well of a Seahorse XFe96 cell culture plate and incubated for 20 min at room temperature. Following the incubation, CellTak was washed out with 200 μl dH_2_O and 250,000 cells were plated per well. The resulting plate was incubated at 37°C in a non-CO_2_ incubator for 45 min. Each drug was diluted in Seahorse media and loaded onto the following ports: Port A, 2 μM Oligomycin (Sigma, 75351); Port B, 2.5 μM FCCP (Sigma, C2920); Port C, 1 μM Rotenone (Sigma, R8875) and 1 μM Antimycin A (Sigma, A8674). Following the incubation, the Mitochondrial Stress Test was performed on a Seahorse Bioanalyzer XFe96 instrument.

### Statistical analysis

The statistical tests and exact group sizes are indicated in each figure legend. Statistical tests were performed in Prism v8.4.1, Excel v16.35, or R v3.5.3. Analysis of differentially expressed genes or accessible regions was performed using edgeR (19) and the Bonferroni FDR correction in R. Differentially expressed genes and differentially accessible regions were determined based on both a false discovery rate (FDR) ≤ 0.05 and absolute log_2_FC ≥ 1. GSEA P-values are determined by permutation testing. Significance between time course qRT-PCR data was determined by two-way ANOVA with Tukey’s post-hoc correction.

## RESULTS

### Memory mice respond with increased magnitude and rapid B cell differentiation kinetics to a secondary influenza challenge

The murine model of influenza infection using heterosubtypic A/PR8/34 (PR8) and A/HK-X31 (X31) viral strains allows for the tracking of memory B cell responses to the shared nucleoprotein (NP) segment of influenza (16, 30, 31). To characterize the differentiation capacity between MBCs and nBs, C57BL/6J mice were infected with a primary dose of PR8 and allowed to rest for 35 days to generate antigen-experienced memory mice. Memory and naïve C57BL/6J mice were then challenged with X31 and the early differentiation kinetics (days 4-7) were monitored by flow cytometry. Memory mice showed enhanced generation of GL7^+^FAS^+^ GC B cells by both frequency and cell number in the spleen and mediastinal draining lymph node (dLN) at all time points assayed (**Fig. 1A, Supplemental Fig. 1**). PC differentiation was low in both naïve and memory mice at day 4, but significantly increased in memory mice compared to naïve at days 5 and 6 post X31 challenge. Naïve mice showed similar frequencies and numbers of CD138^+^ PCs at later time points in the spleen and dLN.

**Figure 1.**
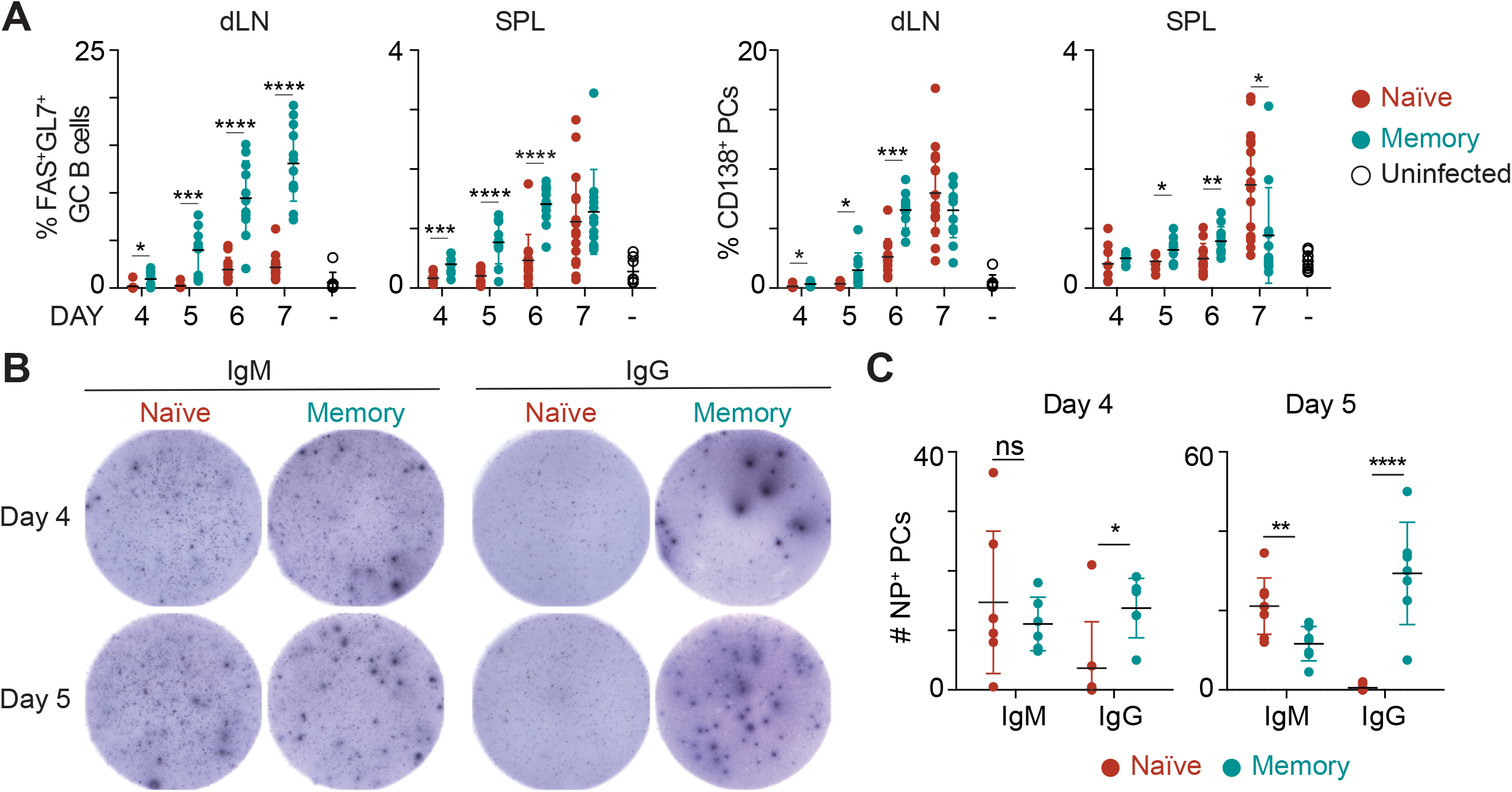
Memory mice show increased kinetics of differentiation and generate more antigen-specific, isotype switched plasma cells. **(A)** Frequency of FAS^+^GL7^+^ germinal center (GC) B cells or CD138^+^ plasma cells (PC) from naïve or memory mice challenged with 0.1 LD_50_ influenza X31 at the indicated time point. Data were collected from the mediastinal draining lymph node (dLN) or spleens (SPL) and are representative of 2 independent experiments with 4-5 mice per group. **(B)** ELISPOT analysis of nucleoprotein (NP)-specific IgM (right) or IgG (left) antibody secreting cells at days 4 or 5 post X31 challenge. Spleens and dLN were pooled, and 500,000 total cells were plated per well. **(C)** Quantitation of IgM and IgG NP-specific ELISPOTS per 500,000 cells. Significance determined using two-tailed Student’s *t*-test. See also **Supplemental Fig. 1**.

To further elucidate differences in influenza-specific PC differentiation, naïve and memory mice were infected with X31 and NP-specific ELISPOTs were performed at days 4 and 5 post infection. NP-specific IgM spots were similar between naïve and memory mice at day 4 post challenge (**Fig. 1B, 1C**). However, there were significantly more NP-specific IgG spots from pooled spleens and dLNs isolated from PR8-primed memory mice than from naïve mice. Thus, as expected from the reports using model antigens and malaria infection (6, 8, 10, 32), the system employed here allows the elucidation of MBCs that form following a viral infection.

### Isotype and influenza-specific MBCs have distinct gene expression profiles

To investigate the molecular properties that underlie enhanced MBC differentiation compared to nBs; at day 35 post PR8 infection, influenza-specific MBCs were isolated using NP B cell tetramers (16). MBCs were subsetted into those that expressed either an IgM or IgG BCR (**Supplemental Fig. 2**). In addition, follicular IgM^+^ nBs from the dLN were isolated from naïve mice and the transcriptome of each population determined by RNA-seq. Principal component analysis (PCA) of all 10,324 detected genes from the RNA-seq data showed that each of the three subsets segregated into discrete clusters (**Fig. 2A**). Principal component 1 (PC1) separated the two MBC populations from the nBs and PC2 distinguished IgM MBCs from the other two subsets. Hierarchical clustering followed by bootstrapping analysis of the differentially expressed genes (DEG; absolute log_2_FC > 1 and FDR < 0.05) revealed that IgM MBCs were a transcriptionally distinct MBC population with both unique and shared gene expression patterns compared with both nBs and IgG MBCs (**Fig. 2B, Supplemental Table 1**). Analysis of the DEG overlap between MBCs and nBs showed 191 signature genes that were similarly expressed in both IgM and IgG MBCs (**Fig. 2C**). Synthetic spike-in mRNAs were used to normalize and quantitate the global mRNA content (22) and revealed a significant increase in IgM and IgG MBCs as compared to nBs (**Fig. 2D**).

To further explore the specific differences between subsets, the relationship between fold change (FC) and significance (FDR) was plotted and gene ontology (GO) (33) analysis was performed on the DEG sets from each comparison group. Both IgM and IgG MBCs showed upregulation of GO terms describing activation, proliferation, and immune system processes, highlighted by *Tbx21* and *Cd80* (**Fig. 2E, 2F**). *Tbx21*, which encodes the transcription factor T-bet, promotes the expression of the chemokine receptor *Cxcr3* (34), which was also upregulated in both MBC subsets, and is required for secondary PC responses to influenza (35). Additional genes that have similar expression patterns in both IgM and IgG MBCs included *Traf1, Vcam1, and Il9r* (Fig. 2G). *Traf1* is a member of the tumor necrosis factor (TNF) receptor family and heterodimerizes with TRAF2 to mediate TNFα-induced NF-κB (36). Similar to *Traf1*, the adhesion molecule *Vcam1* is upregulated in response to NF-κB signaling (37). IL-9R was previously shown to be critical for MBC differentiation from the GC (38). Intriguingly, *Il9r* expression was found to be >3-fold higher in IgM compared to IgG MBCs in our data.

**Figure 2.**
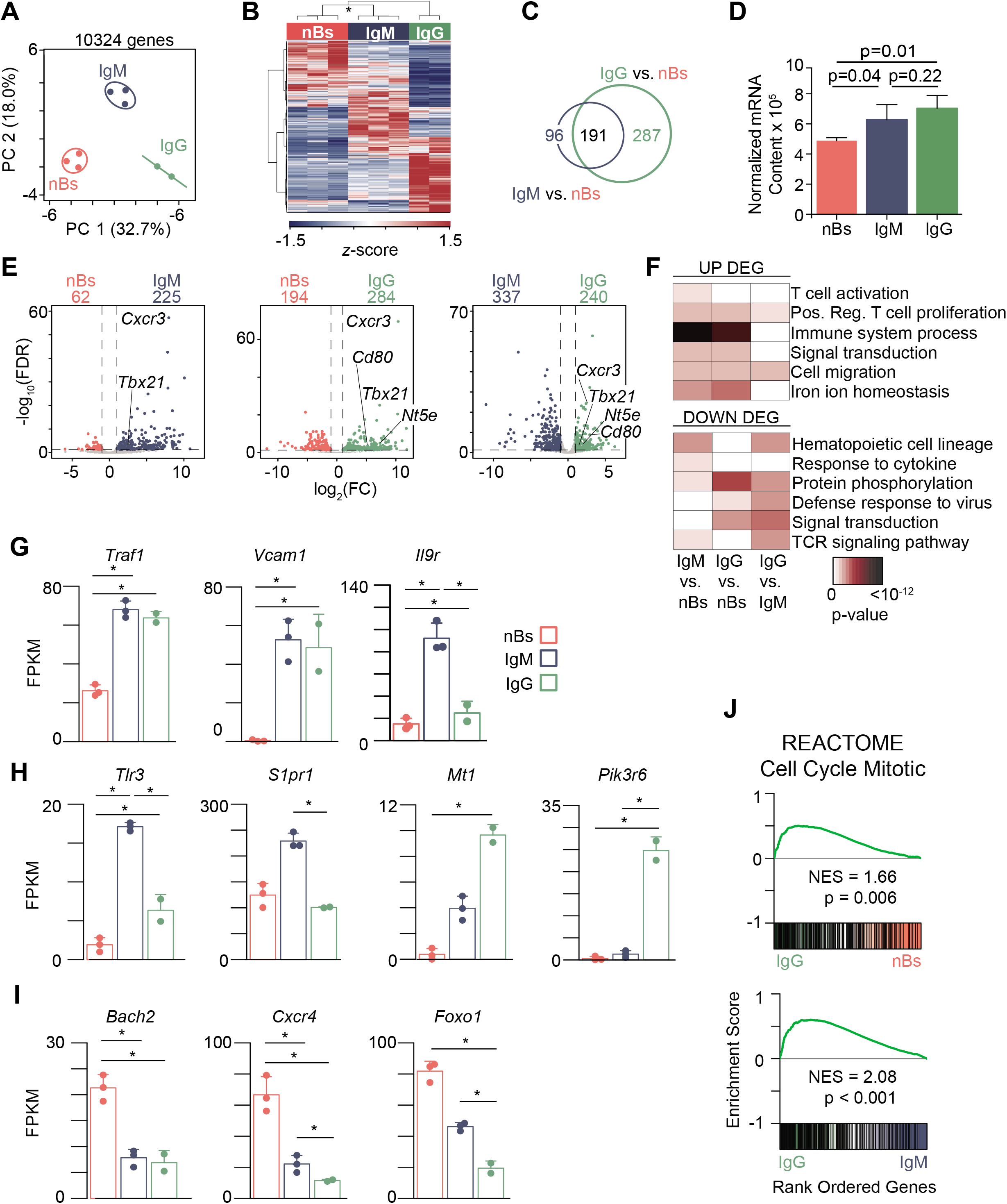
Memory B cells have distinct transcriptional signatures. **(A)** Principal component analysis (PCA) of 10,324 detected genes in follicular IgM^+^ nB (n = 3) from naïve mice and influenza NP-specific IgM (n = 3) and IgG (n = 2) MBC 35 days post-PR8 infection. See also **Supplemental Fig. 2**. **(B)** Heatmap analysis of 901 differentially expressed genes (DEGs). * indicates 100% reproducibility based on bootstrapping analysis. **(C)** Venn diagram showing the overlap of 287 IgM vs. nB DEG (blue circle) with 478 IgG vs. nB DEG (green circle). **(D)** Normalized mRNA content calculated as the sum of all transcripts in a given subset. **(E)** Volcano plots for each comparison with relevant genes upregulated in the MBC subsets indicated. **(F)** Gene ontology (GO) analysis of DEGs in every comparison. Bar plots for key genes highly expressed in: both MBC populations compared to nB **(G)**; IgM or IgG MBC **(H)**; or downregulated in both MBC populations **(I)**. **(J)** Gene set enrichment analysis (GSEA) of the REACTOME Cell Cycle Mitotic gene set in each comparison. NES, Normalized Enrichment Score. * in **G - I** indicates significant differential expression (FDR < 0.05, absolute log_2_FC > 1). See also **Supplemental Fig. 3**.

DEGs uniquely enriched in MBC subsets included *Tlr3* and *S1pr1* in IgM and *Nte5, Mt1*, and *Pik3r6* in IgG MBCs (**Fig. 2H**). Toll-Like Receptor-3, encoded by *Tlr3*, senses intracellular dsRNA and is upregulated in murine B cells upon BCR stimulation (39). S1PR1 is important for egress of lymphocytes from the lymph nodes (40), as well as release of developing B cells from the bone marrow (41). Consistent with its expression on class-switched MBCs (32), *Nt5e*, which encodes CD73, was found to be more highly expressed by IgG MBCs as compared to both nBs and IgM MBCs. *Mt1* plays a role in iron ion homeostasis (42), and *Pik3r6* regulates phosphoinositide 3-kinase signaling, which is activated downstream of BCR triggering (43). The unique expression of *Pik3r6* in IgG MBCs suggests a differential role for this protein downstream of IgM and IgG BCRs (2, 3). Further functional exploration of the DEG using gene set enrichment analysis (GSEA) (44) identified that cell cycle genes were upregulated in IgG MBCs compared to both IgM MBCs and nBs, underscoring their rapid responses to stimulation (**Fig. 2J**). Examples include *Ube2c*, a ubiquitin conjugating enzyme important for degrading target proteins during cell cycle progression (45) and cell cycle genes *Cdc20* and *Mcm10* (**Supplemental Fig. 3A**).

Genes downregulated in both MBC subsets compared to nBs functioned in protein phosphorylation and cellular response to cytokine stimulus (**Fig. 2F**) and included *Bach2, Cxcr4*, and *Foxo1* (**Fig. 2I**). Reduced expression of *Bach2* in MBCs confirms previous observations using model antigens (10). *Cxcr4* is a homing receptor for migration to the bone marrow and between the dark and light zones of the GC. *Foxo1* is critically important for B cell development and maintenance (46). Together, these data indicate that MBCs harbor a gene expression program that is distinct from nBs and sustained following antigen clearance.

### MBCs are epigenetically primed to differentiate to plasma cells

The assay for transposase-accessible chromatin sequencing (ATAC-seq) (47) was performed to interrogate the epigenetic landscape of MBCs and nBs on the same three populations of cells as above. PCA of the 42,231 accessible regions identified revealed that PC1 separated MBCs from nBs and PC2 separated IgM-expressing populations from IgG (**Fig. 3A**). A heatmap of all significant differentially accessible regions (DAR; absolute log_2_FC > 1 and FDR < 0.05) demonstrated a large number of regions in MBC with gains in accessibility compared to nBs (**Fig. 3B, Supplemental Table 2**). Annotation of the chromatin accessibility for 2 kb surrounding all DAR revealed that IgM and IgG MBCs both had overall increased accessibility compared to nBs, with IgG MBCs also displaying a small but significant increase in accessibility over IgM MBCs (**Fig. 3C**). *Zbtb32*, which has been implicated in MBC reactivation (15), showed increased accessibility and a corresponding increase in expression that peaked in IgG MBCs (**Supplemental Fig. 3B**). Additionally, consistent with gene expression changes, DAR were observed surrounding the genes *Tbx21, Nt5e* (CD73), and *Cxcr3*.

**Figure 3.**
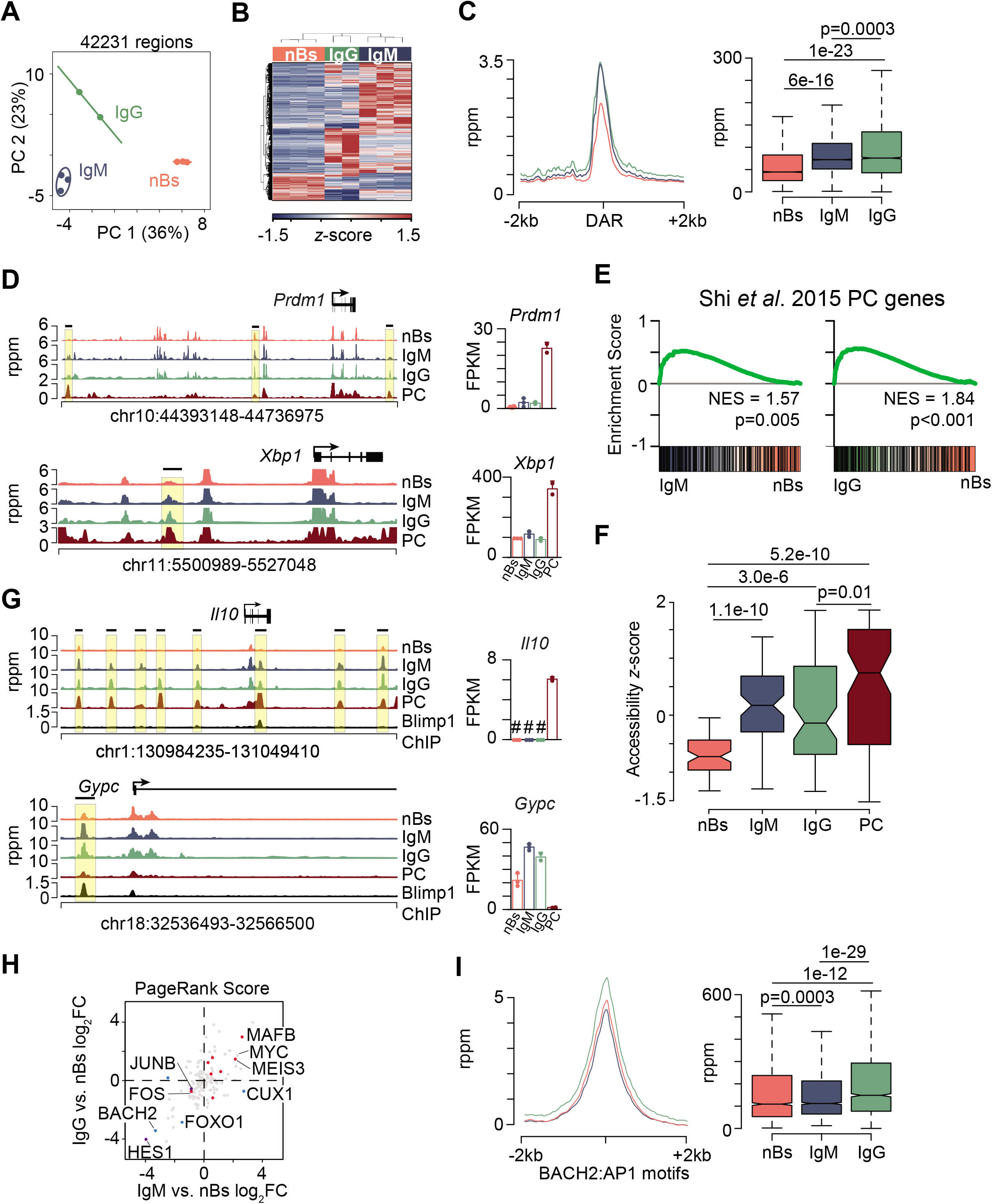
Memory B cells are epigenetically primed to differentiate to plasma cells. **(A)** PCA analysis of 42,231 accessible regions for nB (n = 3) and IgM (n = 3) and IgG (n = 2) MBC defined in **Fig. 2** and **Supplemental Fig. 2**. **(B)** Heatmap showing 6,948 DAR. **(C)** Histogram of all DAR identified in nB, IgM, and IgG MBCs (left) and boxplot quantitating chromatin accessibility (right). **(D)** Genome plots of accessible chromatin (left) and gene expression (right) for select genes. DAR are indicated by a black horizontal bar and yellow box and the upper scale limit was set to show the changes at DAR in MBCs. PC RNA-seq (22) and ATAC-seq (49) datasets were described previously. **(E)** GSEA plots analyzing the Shi *et al*. 2015 PC signature gene set (48) showing enrichment for IgM (right) or IgG (left) MBCs compared to nBs. **(F)** Boxplot of *z*-score normalized accessibility for all DAR that map to the Shi *et al*. 2015 PC genes from **E**. **(G)** Genome plots for indicated gene locus showing accessibility (left) and gene expression (right). BLIMP-1 ChIP-seq data was described previously (50). # denotes genes not detected by RNA-seq for the indicated cell type. DAR are indicated by a black horizontal bar and yellow box and the upper scale limit was set to show the changes at DAR in MBCs. **(H)** Scatterplot of the PageRank fold change for the IgM vs. nB compared to IgG vs. nB. DEG up in nB are indicated in blue, DEG up in IgG only are indicated in purple, and DEG up in both IgM and IgG are indicated in red. **(I)** Histogram (left) and boxplot (right) of accessibility at all BACH2:AP1 binding motifs identified in nB, IgM, and IgG MBCs. Significance for **C, F, I** determined by two-tailed Student’s *t*-test. See also **Supplemental Fig. 3**.

The rapid formation of PCs in memory mice suggested the hypothesis that MBCs might harbor a distinct programming related to PC differentiation. In support of this, DAR were observed that mapped to key PC transcription factors: *Xbp1, Prdm1*, and *Irf4* (**Fig. 3D, Supplemental Fig. 3C**). To examine other genes, a transcription signature derived from genes expressed in PC (48) was used in a GSEA. This analysis revealed that PC genes were significantly enriched in both IgM and IgG MBC populations compared to nB (**Fig. 3E**). The change in chromatin accessibility at DAR mapping to PC signature genes was compared between nBs, MBCs, and previously generated PC data (49). nBs displayed the lowest accessibility with both IgM and IgG MBC subsets exhibiting intermediate levels compared to PC (**Fig. 3F**), indicating that the MBCs may be primed for expression of PC genes. Examples include the plasma cell expressed genes *Prdm1* and *Xbp1*, which displayed regions with higher accessibility in MBCs and little to no accessibility in nBs. While annotated as DEG in our dataset, the overall MBC expression for these genes was low compared to PCs, suggesting chromatin accessibility changes precedes expression changes.

To further examine how the observed accessibility patterns corresponded to PC programming events, BLIMP-1 ChIP-seq data (50) was analyzed along with accessibility data in nBs and MBCs. Both repressed and activated BLIMP-1 target genes displayed a pattern of increased accessibility at the same genomic location that contained a BLIMP-1 binding site and where PC also have open chromatin (**Fig. 3G, Supplemental Fig. 3B**). The BLIMP-1 repressed genes *Gypc* and *Aicda* displayed a binding site and open chromatin at a regulatory site proximal to the promoter. *Il10* and *Tigit* each contained loci that were increased in accessibility in MBC that matched the PC profile. Both of these genes are induced by BLIMP-1 in PC, and despite the MBC accessibility profile, expression was not detected in either nBs or MBCs.

To assess the kinetics of gene expression during reactivation MBCs were enriched along with nB and stimulated to differentiate into PC ex vivo using CD40L, IL4, and IL5. At early time points post stimulation, RNA was extracted and the expression of a set of genes displaying MBC-specific accessible chromatin was assessed by qRT-PCR. Consistent with a role for epigenetic priming, *Zbtb32, Aicda, Il10*, and *Xbp1* were significantly induced more rapidly and to higher levels in stimulated MBCs compared to nB (**Supplemental Fig. 3D**). Importantly, the control gene *Gapdh* was equally induced in both conditions. Thus, these data indicate that MBCs harbor accessibility changes that resemble the PC chromatin architecture and may prime gene expression programs for rapid induction.

To determine additional transcription factor networks that were differentially regulated in MBCs compared to nBs, the PageRank (29) algorithm, which integrates RNA-seq and ATAC-seq data to define transcription factor importance based on changes in target gene expression, was used. For each transcription factor, the fold change in PageRank score between IgG and nB was correlated with the change between IgM and nB to define common changes. Both MYC and MAFB had a high PageRank score in both MBC subsets (**Fig. 3H**). Interestingly, *Myc* was not identified as a DEG in either comparison, despite its target genes and network being activated in MBCs. Consistent with their transcriptional down regulation, BACH2 and FOXO1 were less important in MBC transcriptional networks. In memory CD8 T cells, BACH2 has been shown to repress the activity of AP-1 factors through binding to a similar consensus motif (51). This suggested that lower expression of BACH2 may allow for increased AP-1 activity and therefore accessibility at these motifs. When accessibility was annotated for all instances of the BACH2:AP-1 composite motif in DAR, IgG MBCs displayed significantly increased accessibility compared to both IgM MBCs and nBs (**Fig. 3I**), suggesting there may be a similar mechanism at play in MBCs as in memory CD8 T cells.

### Plasma cell differentiation is enhanced by heme

Intriguingly, both GSEA and GO analysis revealed an enrichment of the gene sets related to iron and heme metabolism as being enriched in MBCs compared to nBs (**Fig. 2F, Fig. 4A**). Examples of iron pathway genes include *Slc40a1, Trf*, and *Hmox1*, which were all significantly upregulated and expressed in both MBC subsets with near background expression in nBs (**Fig. 4B**). Additionally, *Hmox1* displayed promoter chromatin accessibility increases that correlated with its unique expression in MBCs (**Fig. 4C**). HO-1, encoded by *Hmox1*, is a heme-response gene that facilitates its recycling to biliverdin, ferrous iron, and CO (52) and can be used as a readout for intracellular heme levels (53). In agreement with the interpretation that the heme uptake pathway is upregulated in MBCs, paramagnetic bead enriched class switched MBCs displayed significantly higher levels of HO-1 than nBs (**Fig. 4D**), indicating that MBCs retain higher levels of intracellular heme.

**Figure 4.**
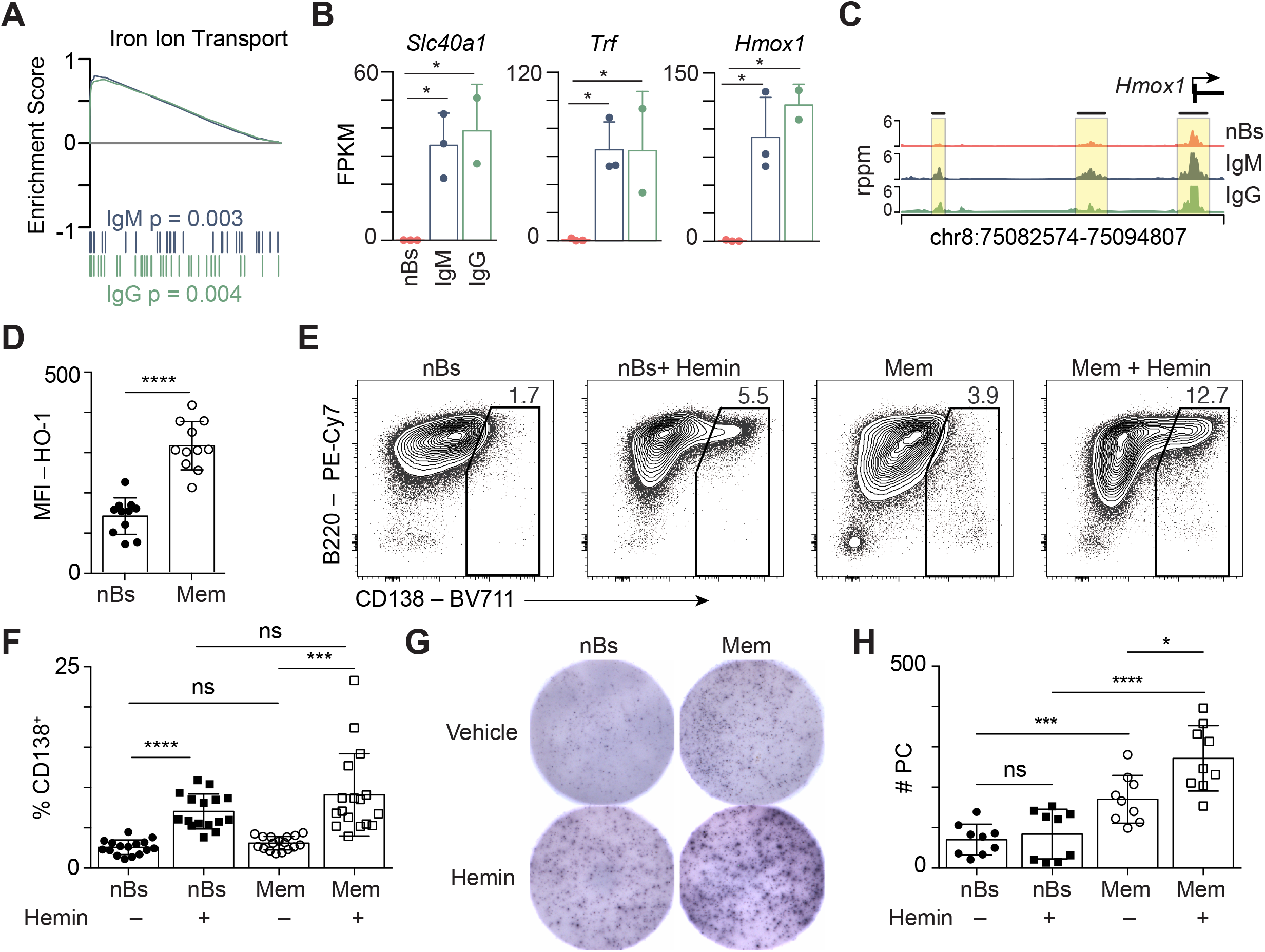
Hemin increases plasma cell formation of murine memory B cells. **(A)** GSEA showing the enrichment of the Iron Ion Transport gene set for IgM (blue) and IgG (green) MBCs compared to nBs. **(B)** Bar plot showing expression of select genes from the Iron Ion Transport gene set in **A**. * indicates significant differential expression (FDR < 0.05, absolute log_2_FC > 1). **(C)** Genome plot of the *Hmox1* locus showing the location of DAR compared to nB. **(D)** Flow cytometry (MFI) quantitation of heme oxygenase-1 (HO-1) in nBs isolated from naïve mice and isotype-switched memory B cells (Mem) that were enriched from PR8-primed mice. **(E)** nBs and Mem were stimulated with CD40L, IL-4, and IL-5 for three days with vehicle or hemin added to the cultures at day 1 as indicated. The formation of CD138^+^ PC was analyzed at day 3 by flow cytometry. **(F)** Quantitation of CD138^+^ PC from **E**. Data are from 3 independent experiments with n = 11. **(G)** Representative ELISPOT wells showing antibody spots per 6250 cells plated from day 3 cultures as in **E. (H)** Quantification of ELISPOT data from **G**. Data are from 3 independent experiments with 3-4 samples per group. Significance for **D, F, H** determined by two-tailed Student’s *t*-test.

To determine if heme modulated PC differentiation, MBCs and nBs were isolated for ex vivo culture and stimulated using CD40L, IL-4, and IL-5 to model T-dependent differentiation conditions. Cultured MBCs differentiated into CD138^+^ PCs to a higher degree than nBs over three days (**Fig. 4E, 4F**). To determine the role of heme in PC differentiation, nBs and enriched MBCs were stimulated in the presence of hemin as above. A 3-fold greater induction of CD138^+^ PCs than matched vehicle treated controls was observed for both nBs and MBCs (Fig. 4E, 4F). Additionally, each culture was subjected to ELISPOT analysis to ensure that hemin treatment produced functional PCs. The addition of hemin significantly enhanced PC formation in MBCs (**Fig. 4G, 4H**). These data indicate that hemin can augment B cell differentiation and enhance the formation of PCs *ex vivo*.

### Epigenetic signatures of MBCs are conserved between mice and humans

To determine if the epigenetic and transcriptional changes observed in mice were also observed in humans, RNA-seq and ATAC-seq data from resting naïve B cells (rN) and class-switched MBCs (SM) from healthy subjects (54) were compare to the data generated here. Of the 10,324 genes expressed in mouse nBs and MBCs, 84% were also detected in the human SM and rN RNA-seq data (**Fig. 5A, pie chart insets**). Reciprocally, 76% of the genes expressed in the human cell types had direct homologues that were detected in the mouse. To determine the concordance of the transcriptional changes, the fold-change of mouse IgG MBCs vs. nBs was plotted against human SM vs. rN B cells. 85 genes were called DEG in both comparisons with 53 having a positive fold change (**Fig. 5A, red**) and 32 having a negative fold change in both species (**Fig. 5A, blue**). Notably, *HMOX1 was* among the conserved upregulated genes along with *ZBTB32, CXCR3*, and *SLC11A1. BACH2, CXCR4*, and *IL21R* highlighted the downregulated conserved genes.

**Figure 5.**
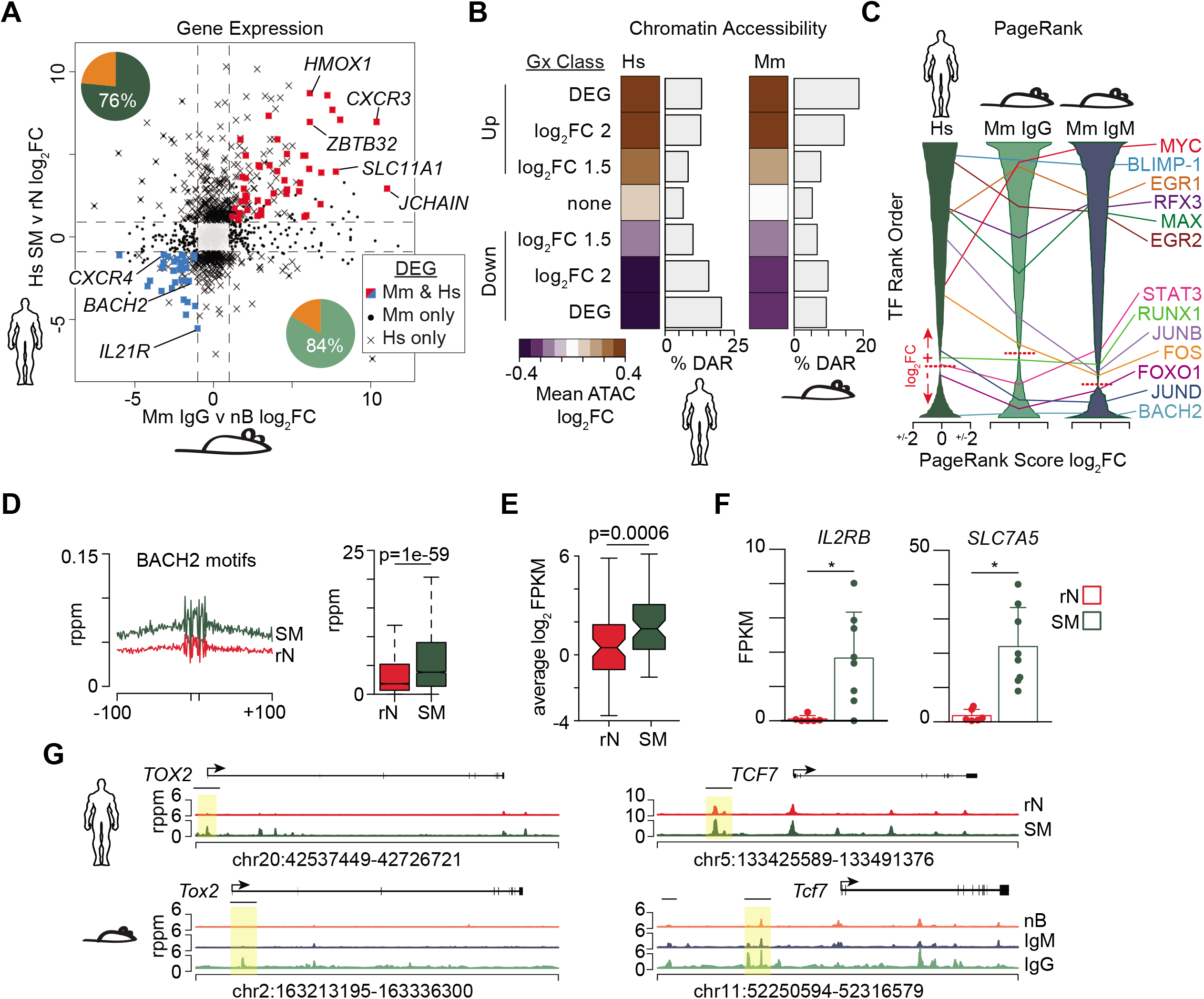
Memory B cell epigenetic signatures are conserved between mouse and human. **(A)** Scatter plot of the fold change for all detected genes between mouse IgG vs. nBs and human isotype-switched memory (SM) vs. resting naïve (rN) reported previously (54). Pie chart indicates the percentage of genes with homologs that were also detected as expressed in the corresponding species. DEG annotations are denoted by symbols shown in the key. **(B)** Heatmap of corresponding chromatin accessibility in human (left) and mouse (right) cell types for peaks surrounding select gene sets. Genes are subsetted based on seven classifications for changes in expression as indicated. The frequency of accessible regions that are annotated as a DAR for each category is shown to the right of each heatmap. **(C)** PageRank analysis (29) was performed on each cell subset and the resulting transcription factors ranked by fold change in memory vs. naïve. X-axis scale indicates PageRank log_2_FC with the dotted line indicating the transition from positive (+) to negative (-) fold changes in the ranked list. Select transcription factors and their location in each list are annotated. **(D)** Histogram (left) and boxplot (right) of accessibility at BACH2 motifs and surrounding sequences. rppm, reads per peak per million. **(E)** Average gene expression of log_2_FPKM normalized data for all BACH2 target genes (55). **(F)** Gene expression bar plots for examples of two BACH2 target genes in rN and SM B cells. * indicates significant differential expression (FDR < 0.05, absolute log_2_FC > 1). **(G)** Genome plots displaying DARs in human (top row) and mouse (bottom row). Highlighted by a yellow box are homologous regions that were identified as DAR. Significance for **D, E** determined by two-tailed Student’s *t*-test.

The global chromatin accessibility profiles between the two species were analyzed by clustering the genes that were conserved between mice and humans into seven transcriptional bins based on fold change. For each bin, the chromatin accessibility fold change between memory and naïve B cells was calculated and averaged for all peaks. A concordance between chromatin accessibility and transcription was observed for genes conserved between the MBCs of humans and mice (**Fig. 5B**). Additionally, up to ~20% of accessible regions were annotated as DAR in the Up and Down DEG bins. Importantly, the genes with no expression changes had the lowest enrichment for accessibility changes.

The PageRank algorithm was applied to the human SM vs. rN comparison and the mouse IgG and IgM MBCs versus nBs. Each transcription factor was ranked from the most important in MBCs to nBs. Many transcription factors showed a similar ranking of importance in each of the three comparisons (**Fig. 5C**). Among those that had high importance in MBCs included MYC, BLIMP-1, EGR1, RFX3, MAX, and EGR2. Among those that had high importance in nBs were BACH2, FOXO1, and JUND (**Fig. 5C**). To further explore the change of BACH2 transcriptional networks in MBCs, all peaks were annotated for the presence of a BACH2 binding site motif and the chromatin accessibility in SM and rN calculated (**Fig. 5D**). We observed significantly more accessibility at BACH2 binding motifs in human SM compared to rN. Additionally, the overall expression of genes that were predicted to be BACH2 targets (55) was increased in SM cells (**Fig. 5E**). For example, the BACH2 targets, *IL2RB*, a component of the IL-2 receptor, and *SLC7A5*, a subunit of the neutral amino acid transporter CD98, were both increased in SM (**Fig. 5F**). Because BACH2 is described as a transcriptional repressor and its expression is decreased in MBCs, this suggests that the BACH2 binding sites, which are also shared by other members of the AP-1 family of factors, are derepressed.

Examples of regions that are directly conserved between mice and humans and have similar changes in programming include *Tox2* and *Tcf7* (**Fig. 5G**). Both of these genes display DAR that have increased accessibility in MBCs compared to naïve cells at syntenic genomic regions. *Tox2* drives T follicular helper cell differentiation (56) and is regulated by BACH2 in B cells (55). *Tcf7* encodes the gene TCF1 that is required for CD8 memory formation and marks resource stem cells that are capable of reseeding the effector population (57, 58). Thus, human and mouse MBCs share a conserved programming with respect to chromatin accessibility and gene expression.

### Hemin enhances plasma cell differentiation of human memory B cells

Consistent with mouse MBCs, the Iron Ion Homeostasis gene set was significantly enriched in MBCs from humans (**Fig. 6A**). Examples of these genes include *HMOX1, HFE*, and *SLC11A1* (**Fig. 6B**). HO-1 expression was used as a readout for intracellular heme levels (53) and, as in the mouse, SM cells displayed increased expression of HO-1 compared to rN cells (**Fig. 6C**). The role of heme in MBC reactivation was evaluated by stimulating SM cells with a cytokine cocktail that promotes PC differentiation over six days (59) in the presence of vehicle or hemin. The MBC cultures that were treated with hemin exhibited increased formation of CD38^+^CD27^+^ PC (Fig. 6D, 6E). Additionally, the formation of a more mature population of PC that were CD38^+^CD138^+^ (60) was increased in the presence of hemin (**Fig. 6F, 6G**). ELISPOT analyses of these cultures confirm increased antibody secretion in cultures treated with hemin (**Fig. 6H**). Thus, hemin directly influences the *ex vivo* generation of plasma cells from human MBCs.

**Figure 6.**
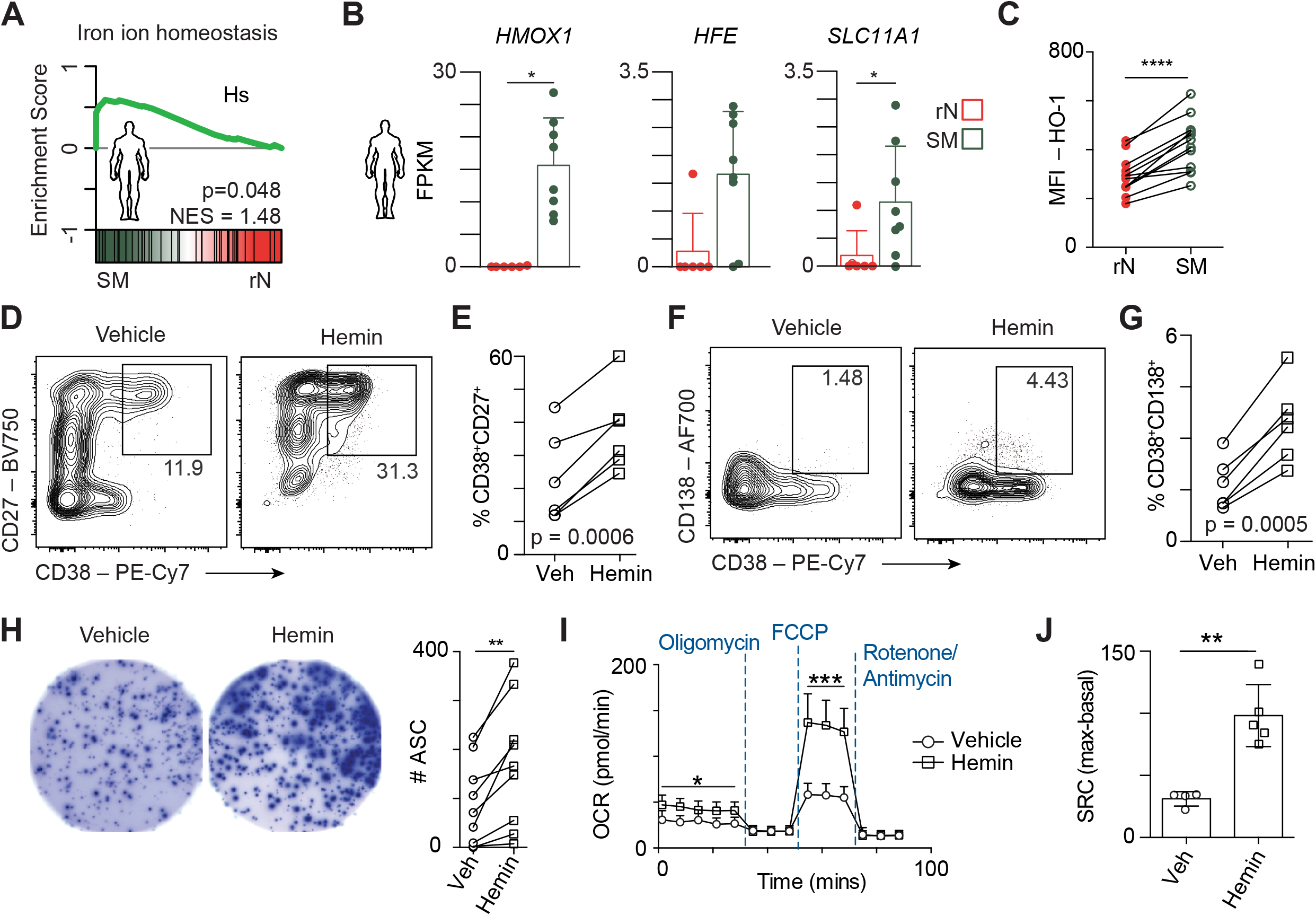
Hemin augments human plasma cell differentiation and mitochondrial metabolism. **(A)** GSEA for the Iron Ion Homeostasis gene set between human SM vs. rN B cells. **(B)** Bar plots showing gene expression for the indicated genes from the Iron Ion Homeostasis gene set in **A**. * indicates significant differential expression (FDR < 0.05, absolute log_2_FC > 1). **(C)** MFI quantitation of HO-1 in rN and SM cells isolated from total PBMCs from 13 human healthy donors. **(D)** Representative flow cytometry plots of SM from a single donor stimulated with R848, CD40L, IL-21, IL-2, BAFF, and human insulin as described previously (59) and supplemented with vehicle control or hemin showing the formation of CD38^+^CD27^+^ PC. **(E)** Quantitation of CD38^+^CD27^+^ PC from 6 individual donors differentiated as in **D. (F)** Representative flow cytometry of CD38^+^CD138^+^ PC from cultures differentiated as in **D. (G)** Quantitation of CD38^+^CD138^+^ PC from 6 individual donors from **F. (H)** Representative ELISPOT showing PC antibody spots from 3125 cells plated from the cultures described in **D** (left) and quantitation of spots from 9 individual donors (right). **(I)** Memory B cells were differentiated as in **D** then harvested and washed in Seahorse media before being subjected to a Mitochondrial Stress Test on a Seahorse Bioanalyzer. **(J)** Spare respiratory capacity was quantified as the difference between first measurement following FCCP injection and the last measurement before oligomycin injection and shown as a bar plot. Data are representative of five patients from two independent experiments. Significance in **C, E, G, H, J** determined by two-tailed Student’s *t*-test.

### Hemin treatment augments mitochondrial metabolism

The mitochondria is the primary place that iron resides within cells where it is integral to the functions of the electron transport chain and the ability of cells to perform oxidative phosphorylation (52). Previous observations showing that oxidative phosphorylation promotes PC differentiation (61) suggested that the increased PC differentiation in the presence of heme may be accompanied by metabolic changes. To test this, SM B cells were enriched from human PBMCs, stimulated to differentiate (59) in the presence or absence of hemin as above, and the resulting cultures were analyzed for their usage of oxidative phosphorylation by extracellular flux assay on a Seahorse bioanalyzer. Hemin treated samples showed significantly higher basal and maximal respiration (**Fig. 6I**), which also resulted in increased spare respiratory capacity (**Fig. 6J**). These data indicate that hemin facilitates increased mitochondrial metabolism and oxidative phosphorylation *ex vivo*.

## DISCUSSION

To distinguish the epigenetic and transcriptional properties contributing to B cell memory, the chromatin accessibility and transcriptome profiles of influenza-specific IgM and IgG MBCs were determined and compared to those of nBs. An overall increase in MBC accessibility was identified that correlated with PC programming and indicated that prior B cell activation imparted an epigenetic state that was distinct from antigen-inexperienced nBs. IgM and IgG MBCs were also distinct from each other, agreeing with previous observations that each MBC subset may be predisposed for different reactivation fates (62). Importantly, MBC programming was largely conserved between mice and humans with the heme and iron homeostasis pathways significantly upregulated in both species. Increased MBC heme content was validated and ex vivo differentiation in the presence of hemin enhanced PC differentiation. Together, these data identify an epigenetic programming that is unique to MBCs and highlight mechanisms for enhanced recall responses upon secondary antigen exposure.

The chromatin accessibility data provide evidence for a primed epigenetic architecture as one mechanism that allows for more rapid responses for MBCs compared to nBs. The maintenance of accessible chromatin at specific loci in MBCs may act to reduce the number of reprogramming steps necessary for many functions, including the formation of GC and PC following reactivation. Indeed, MBC displayed higher expression of genes related to the cell cycle, B cell activation, signal transduction, and migration. Many of these genes showed a graded expression that was high in IgM and maximal in IgG MBCs compared to nBs. Of particular interest was the enhanced expression and accessibility surrounding genes unique to PC in MBCs. PageRank analysis of both mouse and human MBCs revealed BLIMP-1 as an enriched regulator of the MBC transcriptional network, indicating that aspects of the PC program is partially engaged or primed in MBCs. In support of this, the 3D architecture of the *Prdm1* locus, which encodes BLIMP-1, is reorganized in activated B cells prior to *Prdm1* expression in PC (63). Chromatin accessibility priming was also observed in memory CD8 T cells and helped facilitate rapid induction of effector responses upon secondary stimulation (64). Similar to memory CD8 T cells, a set of primed genes demonstrated rapid induction and higher expression in MBC compared to nB. Therefore, in a similar manner, MBCs maintain an accessibility programming that serves as a molecular signature of prior activation and may improve the fidelity and speed of differentiation during secondary antigen encounters.

The presence of differentially accessible chromatin implies that transcription factor activity is different between MBCs and nBs. Consistent with previous findings (10), both human and mouse MBCs displayed lower expression of *Bach2*. BACH2 functions by partnering with various AP-1 factors and represses its target genes (51). Both species displayed increased accessibility at BACH2/AP-1 binding motifs. Intriguingly, such target genes such as *Prdm1* (mouse) and *SLC7A5* and *IL2RB* (human) were increased in expression, indicating BACH2 activity was reduced in MBCs and that other AP-1 family members were dictating the change in expression. This was supported by PageRank analysis, which found both MYC and a set of AP-1 factors to be more important in MBCs and might allow more rapid induction of metabolic changes, proliferation, and differentiation to PC. Altered MBC metabolic states were indicated by enhanced mRNA and heme content compared to nBs. Additionally, activation of the AKT pathway, which regulates cellular metabolism through mTOR was indicated by reduced expression and network importance of *Foxo1* (65). Furthermore, *Zbtb32*, which acts to restrict secondary responses (15), was upregulated in IgM and was maximally expressed in IgG MBC. Thus, altered MBC transcription factor networks provide mechanisms to rapidly modify metabolism, differentiate to PC, and ultimately regulate humoral responses.

One of the most striking conserved differences between MBCs and nBs was the increased expression of heme biosynthetic genes in MBCs. Heme enhanced the formation of PC from both human and mouse MBCs during ex vivo differentiation. Heme is synthesized and primarily found in the mitochondria where it is an essential component of cytochrome proteins that function in the electron transport chain during oxidative phosphorylation (52). BACH2 and BACH1 both negatively regulate heme biosynthesis in part through direct binding to the *Hmox1* promoter (55, 66). Heme can also directly bind BACH2 leading to its degradation and derepression of the heme biosynthesis pathway (53). Heme binding is not the only mechanism controlling BACH2 activity, as the PI3K pathway can also lead to phosphorylation of and decreased nuclear localization of BACH2 (67). Given that heme is an essential component of the electron transport chain and our previous findings linking PC differentiation to oxidative phosphorylation (61), this relationship was explored ex vivo. Differentiation in the presence of heme resulted in increased oxidative phosphorylation. In T cells iron is important for activation induced proliferation and mitochondrial function (68). Thus, by upregulation of the components to import and generate heme, MBCs are primed to rapidly increase their metabolism towards modes that promote PC differentiation and support the demands of antibody secretion.

The importance of heme and iron biogenesis for MBCs is also observed in many human disorders. Patients with combined immunodeficiency caused by a missense mutation in *TFRC*, the transferrin receptor, have reduced MBC frequencies as a proportion of total B cells (69). TFRC is required for iron uptake and this observation suggests that iron is important in either the formation or maintenance of MBCs. In multiple myeloma, the iron exporter ferroportin-1 (*SLC40A1*) is decreased, resulting in increased cell-associated iron and poor prognosis (70, 71). Consistent with previous reports (53), administration of heme to mice had no measurable effect on the immune response to influenza (data not shown). However, mice deficient in HO-1, which degrades heme, have increased serum antibody titers (72), indicating heme regulates B cell function in vivo. Thus, targeted modulation of iron metabolism and biogenesis may be a mechanism to promote antibody responses, as well as limit PC differentiation and cell survival in cancer and potentially autoimmune settings. In summary, these data define a conserved epigenetic state centered around primed chromatin accessibility in MBC that provides a molecular history of prior activation and likely contributes to enhanced functions and responses upon secondary challenge.

## Supporting information

Differentially expressed genes between IgM and IgG MBC and nBs.

Differentially accessible regions between IgM and IgG MBC and nBs

## ACKNOWLEDGMENTS

We thank R. Butler for animal husbandry and care; R. Karaffa for cell sorting at the Emory Flow Cytometry Core; and Drs. E. Zumaquero and F.E. Lund (University of Alabama, Birmingham) for guidance with human ex vivo MBC cultures and differentiation.

## ABBREVIATIONS

ATAC-seq: Assay for transposase accessible sequencing;
DAR: differentially accessible region;
DEG: differentially expressed gene;
dLN: draining lymph node;
FC: fold-change;
FDR: false-discovery rate;
GO: gene ontology;
SEA: gene set enrichment analysis;
MBCs: memory B cells;
nBs: naïve B cells;
NP: nucleoprotein;
rN: resting naïve B cells;
PC: principal component;
PCA: principal component analysis;
PCs: plasma cells;
SM: class-switched memory B cells

## SUPPLEMENTAL FIGURES

**Supplemental Figure 1.**
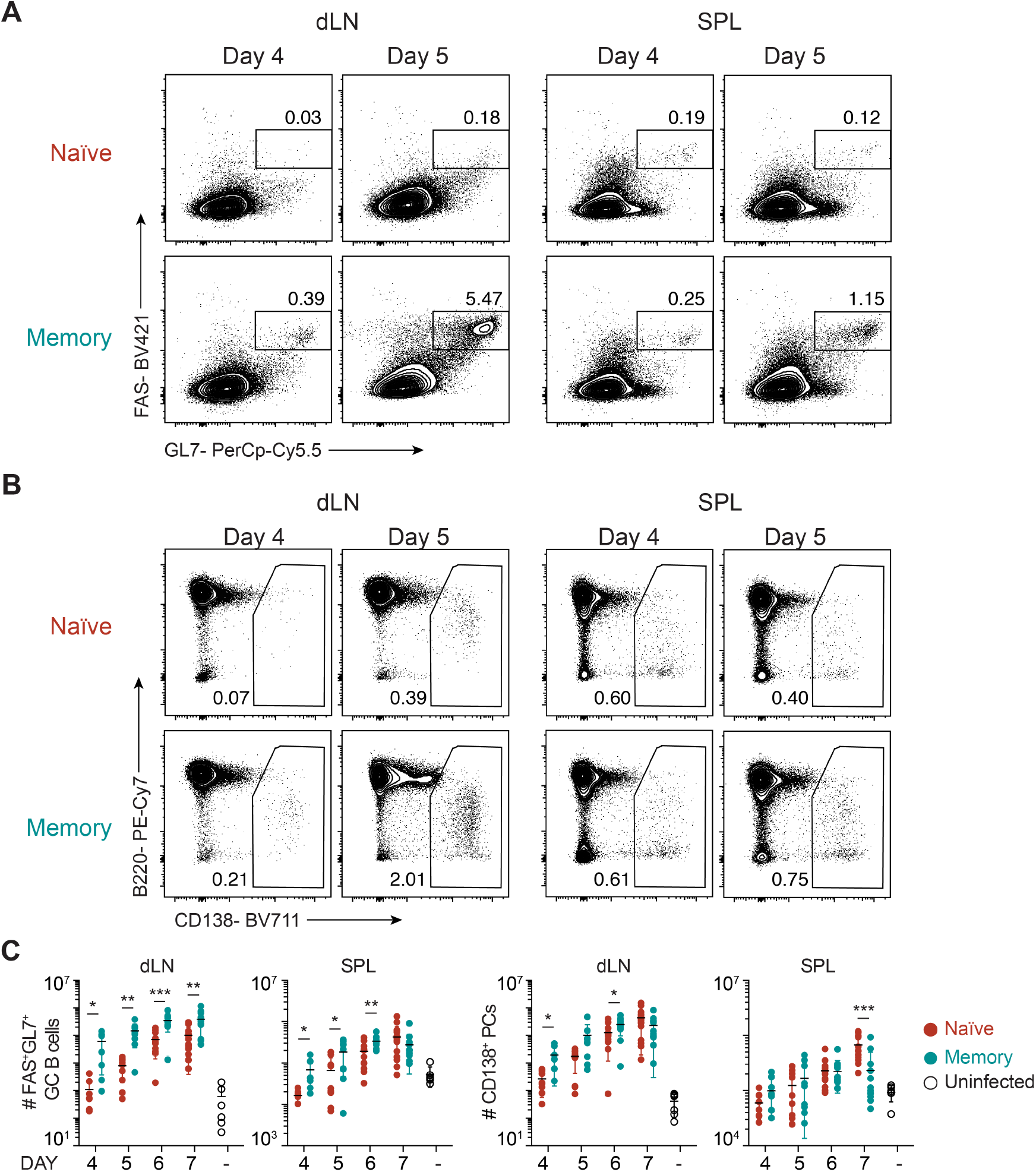
B cell differentiation kinetics of naïve and memory mice. Representative flow cytometry showing the percentage of **(A)** FAS^+^GL7^+^ germinal center (GC) B cells and **(B)** CD138^+^ PCs for the dLN (left) and spleen (SPL, right) for the indicated day in naïve or memory mice. **(C)** Quantitation of the absolute number of GC B cells and PCs from **A** for the indicated day. Significance determined by two-tailed Student’s *t*-test. Data related to **Fig. 1**.

**Supplemental Figure 2.**
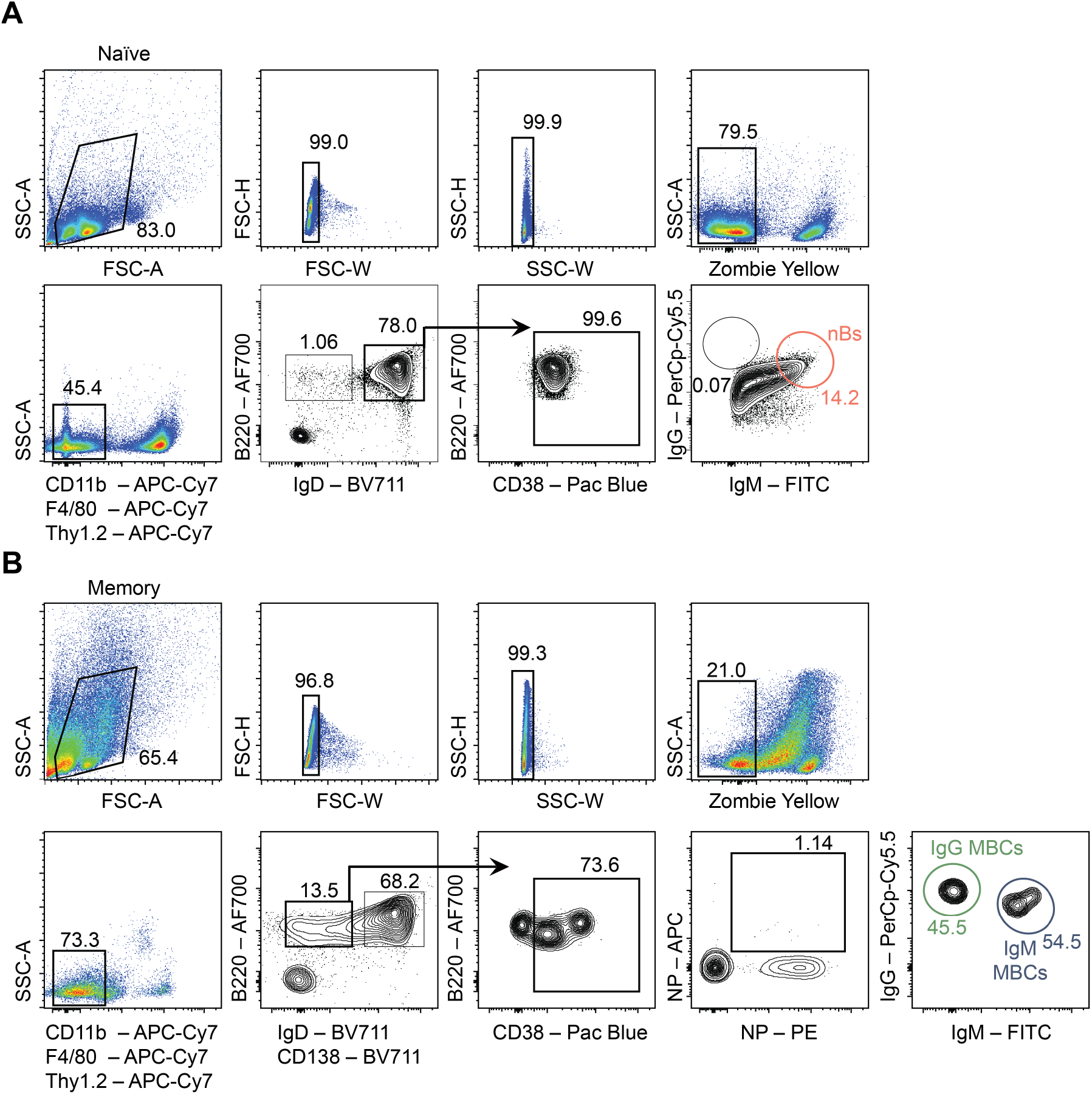
FACS isolation of influenza-specific MBC and nBs. Gating strategy showing the FACS isolation of **(A)** nBs from the mediastinal LN and **(B)** IgM and IgG MBCs from pooled spleens and mediastinal LNs from 4 mice per sample. Data related to **Fig. 2** and **Fig. 3**.

**Supplemental Figure 3.**
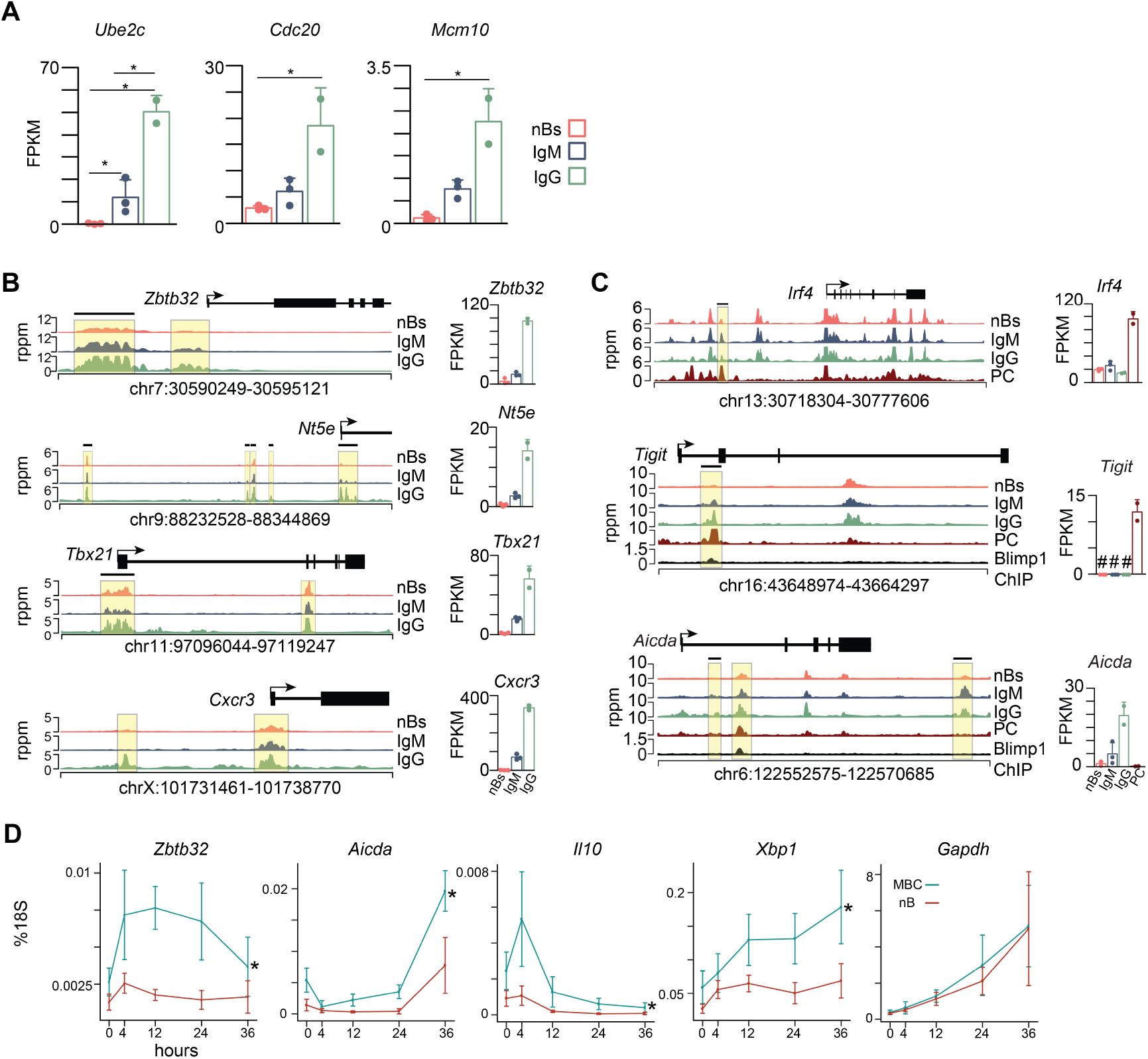
MBCs harbor unique gene expression and accessibility programs. **(A)** Example bar plots of genes that are differentially expressed in MBCs compared to nBs. **(B)** Genome plots (left) and gene expression (right) for select DEG containing corresponding DAR between MBC subsets and nBs. DAR are indicated by a black horizontal bar and regions of interest by a yellow-shaded box. **(C)** Genome plots (left) and gene expression (right) for PC signature genes or genes regulated by BLIMP-1 in PC. # denotes genes not detected by RNA-seq for the indicated cell type. DAR are indicated as above. PC RNA-seq (22), ATAC-seq (49), and BLIMP-1 ChIP-seq (50) datasets were described previously. **(D)** Quantitative RT-PCR time course for the indicated gene in nB or MBC stimulated with CD40L, IL4, and IL5. Data are plotted as a percentage of 18S rRNA with error bars representing SD. Each time point represents cells from 6-9 mice from two independent experiments. Statistical difference was determined by twoway ANOVA with Tukey’s post-hoc correction with * indicating P < 0.05. Data related to **Fig. 2** and **Fig. 3**.

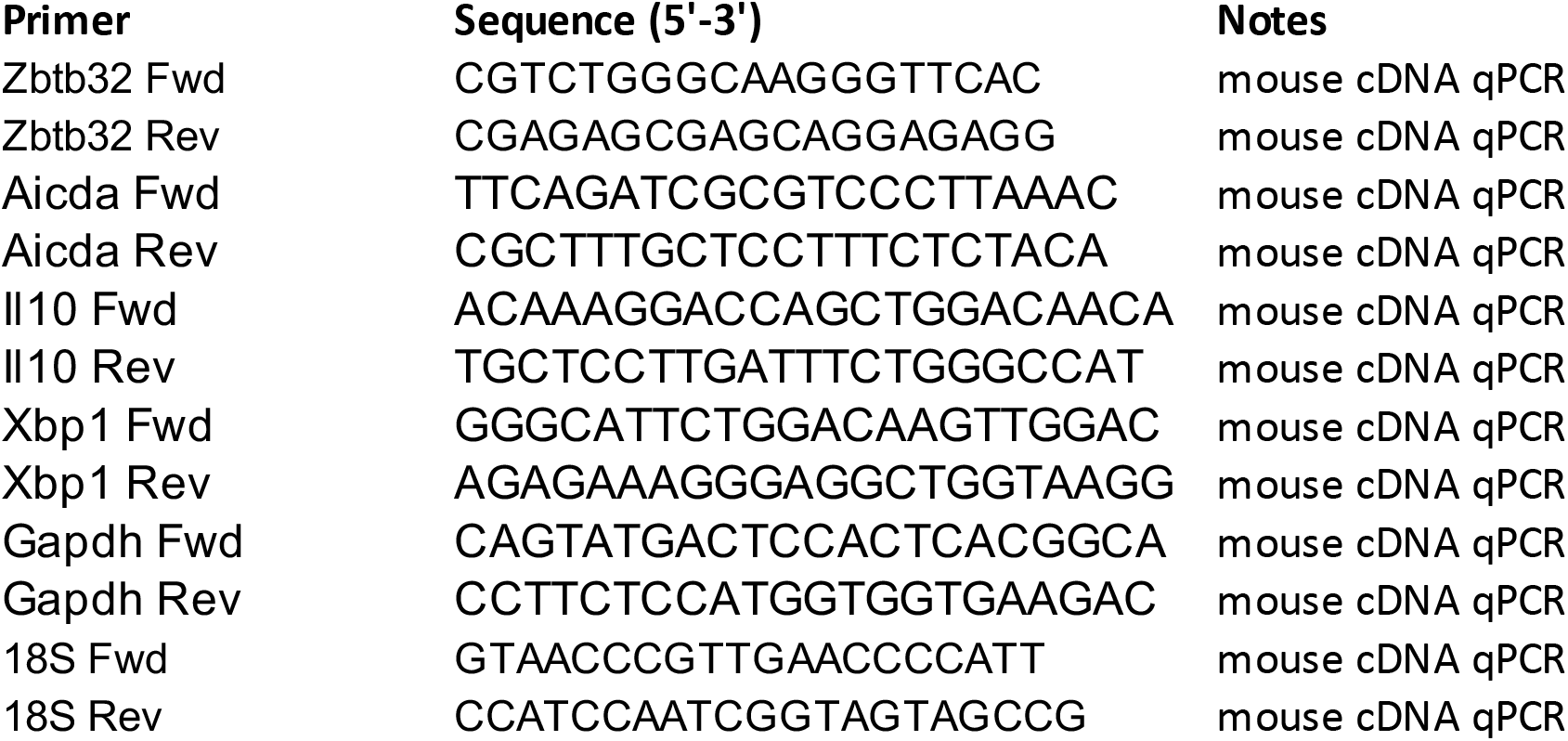

